# Dynamic Estimation of Spatially Interactive Networks (DESINE) Reveals Constrained Brain Repertoire in Schizophrenia Linked to Clinical and Cognitive Symptoms

**DOI:** 10.64898/2026.05.20.726604

**Authors:** Krishna Pusuluri, Godfrey Pearlson, Armin Iraji, Vince D. Calhoun

**Author notes:** Corresponding author: Research Neuroscientist.

## Abstract

**Background:** While resting-state fMRI demonstrated that brain networks are spatially dynamic (expanding, shrinking, and changing complexity over time), understanding the transient spatial network interactions that remain poorly characterized is critical for revealing the mechanisms underlying brain disorders.

**Methods:** We introduce DESINE (Dynamic Estimation of Spatially Interactive Networks), a novel framework using joint density distributions (2D histograms) of voxel-wise activity to quantify 4D spatial network interactions across sliding windows. We analysed transient deviations from the average functional state using root-mean-square error (RMSE) and mean absolute deviation (MAD), and characterized recurring interaction patterns using k-means clustering. We applied DESINE to 91 network pairs (14 networks) in a cohort of 508 subjects (315 healthy controls; 193 patients with schizophrenia, SZ).

**Results:** SZ is characterized by a significantly “constrained dynamic repertoire” of network interactions. SZ patients showed markedly lower means and standard deviations for both RMSE and MAD metrics across network pairs, particularly in regions of high activity, indicating systematic rigidity. Cluster analysis revealed significant alterations in state affinity metrics, suggesting a global breakdown in the brain’s capacity to preserve diverse, high-fidelity spatial configurations. Critically, these interaction metrics were associated with cognitive performance, symptom scores on the positive and negative syndrome scale, and chlorpromazine equivalent drug scores.

**Conclusions:** This work introduces DESINE as a global, voxel-agnostic framework for characterizing time-varying spatial interactions. Our findings highlight spatial rigidity as a fundamental feature of psychopathology, suggesting that the inability to express a diverse range of spatial interactions is a factor underlying cognitive deficits in schizophrenia.

## 1. Introduction

The investigation of brain dynamics in resting-state functional magnetic resonance imaging (rsfMRI) traditionally centered on the “temporal chronnectome” that studied time-varying connectivity between spatially fixed regions or networks [1-4]. While this approach has revealed critical insights into the dysconnectivity characteristic of schizophrenia (SZ) [2,5], it relies on the implicit assumption that the spatial boundaries of functional networks are fixed over time. Recent evidence challenged this paradigm, demonstrating that intrinsic connectivity networks (ICNs) are inherently dynamic, exhibiting continuous voxel-wise expansion, contraction, and structural reorganization over time [6-10].

We previously introduced the concept of the 4D dynamic spatial brain networks with (3D) voxel wise changes over time and quantified the temporal changes in network volumes, revealing significant “volumetric coupling” where networks grow and shrink in a synchronized fashion [8]. We found that SZ is associated with reduced volumetric variability and lower coupling between networks such as the cerebellar and subcortical circuits. Our work integrating spatial complexity into this framework [9] showed that networks in SZ are often “stuck” in disorganized, irregular spatial configurations with diminished “dynamic flexibility”, failing to fluidly change structural complexity to meet cognitive demands.

However, while we characterized the dynamics of individual networks, the question of how these networks interact spatially remains an open frontier. Functional integration and segregation are not merely temporal events; they are governed by the transient spatial overlap and receding boundaries of voxel-level activity distributions [6,10]. We previously studied network interactions through simple correlations between time courses of spatial changes in volume or complexity [8,9], but the nuances of how the joint density distributions of two networks evolve are not fully captured. For instance, two networks might maintain the same average correlation while their spatial boundaries overlap in vastly different configurations.

In this study, we introduce DESINE (Dynamic Estimation of Spatially Interactive Networks) as a global, voxel-agnostic framework to quantify 4D spatial brain network interactions using joint density distributions (2D histograms) [11] that quantify the frequency of various paired combinations of voxel-level intensities or activities across two networks. By generating 2D histograms for network pairs across sliding windows, we can capture the magnitude of deviation from the group mean functional state using Root-Mean-Square Error (RMSE) and Mean Absolute Deviation (MAD). Applying k-means clustering, we characterize the recurring state space of spatial interactions and quantify subject-level affinity for these states using similarity and distance metrics.

We hypothesize that the “spatial rigidity” and “maladaptive complexity” previously observed in individual networks in SZ will manifest as a constrained dynamic repertoire of inter-network interactions [8,9]. Specifically, we expect SZ patients to exhibit lower magnitudes of transient spatial deviations and a loss of fidelity to healthy interaction states. We seek to link these interaction-level disruptions to clinical symptom severity (Positive and Negative Syndrome Scale - PANSS) [5,12], Chlorpromazine (CPZ) equivalent drug intake [13,14], and cognitive performance (processing speed, working memory, verbal/visual learning) [15], providing a mechanistic link between spatial interaction deficits and the clinical phenotype of schizophrenia [8].

## 2. Methods

### 2.1 Data Collection and Preprocessing

We analyzed 3-Tesla rsfMRI data from 508 subjects (315 controls, CN; 193 SZ patients) across three datasets: Functional Imaging Biomedical Informatics Research Network (FBIRN) [2], Center for Biomedical Research Excellence (COBRE) [16], and Maryland Psychiatric Research Center (MPRC) [17], approved by respective institutional review boards. Preprocessing performed using the statistical parametric mapping toolbox (SPM12) included several steps to prepare the magnetization data. The initial five volumes were excluded to ensure magnetization equilibrium. Rigid body motion correction was applied to address head movement, followed by slice-timing correction. Subject data was aligned to the Montreal Neurological Institute (MNI) template and resampled to isotropic voxels of 3 mm^3^. Spatial smoothing was performed using a 6-mm full width at half maximum Gaussian kernel. To mitigate noise and nuisance signals, the voxel-level time courses underwent detrending, despiking, motion correction, and filtering. More information regarding the demographics, datasets, and preprocessing is available in our prior work [10].

### 2.2 The DESINE Framework

Our DESINE framework analyzes time-varying spatial interactions between intrinsic connectivity networks. Unlike traditional methods focused on temporal synchronization, DESINE quantifies voxel-level co-evolution of network topographies through the following stages (Figure 1):

1. **Group-level sICA:** Group-level spatially constrained independent component analysis (sICA) was performed on rsfMRI data using Group ICA of fMRI Toolbox (GIFT) [4], employing a model order of 20 components consistent with prior research [6,8,9]. Of the 20 identified components, 14 were determined to be neuro-relevant brain intrinsic connectivity networks (ICNs) (Figure.2) based on their spatiotemporal properties [10]. The sICA procedure successfully identified and separated these 14 ICNs along with their associated time courses. These networks are: VIS-P (visual primary), SUB (subcortical), MTR-P (somatomotor primary), CER (cerebellar), ATN (attention-dorsal), FRNT (frontal), MTR-S (somatomotor secondary), FPN-R (frontoparietal right), VIS-S (visual secondary), pDMN (posterior default mode), FPN-L (frontoparietal left), SN (salience), TEMP (temporal), and aDMN (anterior default mode).
2. **Subject-level MOO-ICAR:** The second step utilized a sliding-window approach combined with sICA for each subject across time windows [10]. This was achieved using multi-objective optimization ICA with reference (MOO-ICAR) [18,19]. By using the components from group-level ICA analysis (previous step) as references, MOO-ICAR ensured that ICNs corresponded across both subjects and time windows. Our previous work demonstrated MOO-ICAR’s effectiveness in estimating large-scale networks from short time segments [20]. This process allowed us to capture, for each subject, ICNs that varied spatially over time windows, along with their respective time courses within each window. A sliding window length of 60 s (30×TR, with a repetition time TR of 2s) was chosen, consistent with prior research [6] and falling within ranges recommended by multiple studies [4]. Further specifics regarding the first two steps of the analysis pipeline are available in [10].
3. **Interaction Analysis (2D Histograms):** The third step involved calculating joint density distributions (2D histograms) for each subject at every sliding window for a specific brain network pair. This employed normalized (z-scored) voxel-level activity for both the networks, ranging from -4 to +4 (mean at 0) in steps of size 0.1, resulting in a 2D histogram of size 81×81. To isolate differential dynamics rather than overall activity, the mean 2D histogram calculated across all subjects and windows was subtracted from window-level 2D histograms. All further analysis was performed on these 2D difference histograms. This process was executed for 91 possible pairs of 14 ICNs. A few examples of 2D differences histograms and corresponding network activity maps for the network pair VIS-P×FPN-R are shown in Figure.3 [11].
4. **2D Histogram Metric Extraction:** We performed detailed analyses on window-level 2D difference histograms for each network pair and calculated several subject-wise metrics.
  a. **Scalar Metrics:** Root-mean square error (RMSE) and mean absolute deviation (MAD) were calculated for each window to quantify the magnitude of deviation from the mean state.

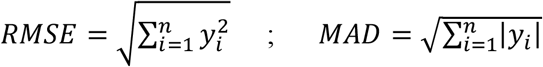 Here, *y*_*i*_ represents each pixel of the 2D difference histogram and ‘n’ represents the total number of pixels (n = 81 × 81). RMSE quantifies the overall magnitude of the differences from mean 2D histogram, penalizing larger deviations more, while MAD measures the average absolute difference from the mean, offering a more robust measure against outliers. We calculate the mean and standard deviation (std.) of these scalar RMSE and MAD values across windows for each subject. These metrics measure the magnitude of the transient spatial interaction dynamics relative to the group average for each network pair. Higher means imply that the subject’s typical interaction state is far from the average, suggesting high overall dynamic range. Higher std. implies the subject exhibits a wider range of dynamic states, fluctuating between very high and low deviations, suggesting high dynamic flexibility and instability.
  b. **Scalar Metrics - Quadrant Analysis:** We extend the scalar metric analysis to specific regions of the histogram and calculate MAD/RMSE metrics and their mean/std. for each of the four quadrants. This approach is valuable because a single, overall metric might obscure a significant difference that is concentrated in just one of these quadrants. It allows us to pinpoint exactly where in the joint activity space of two networks the group differences lie.
    i. Quadrant I (Network 1 > 0, Network 2 > 0): Captures interactions when both networks are active.
    ii. Quadrant II (Network 1 < 0, Network 2 > 0): Captures interactions when network 1 is inactive and network 2 is active.
    iii. Quadrant III (Network 1 < 0, Network 2 < 0): Captures interactions when both networks are inactive.
    iv. Quadrant IV (Network 1 > 0, Network 2 < 0): Captures interactions when network 1 is active and network 2 is inactive.
  c. **K-means Clustering:** To analyze the 2D difference histograms for each network pair, we employed the k-means algorithm to cluster data across subjects and windows. Determining the optimal number ‘k’ of clusters was achieved using the elbow criterion on a smaller subsample of 50 subjects. Specifically, k-means clustering was run for k values ranging from 2 to 50, repeated for robustness with 5 distinct random initialization points for the cluster centroids and up to a maximum of 1000 iterations. The value k=35 was selected as optimal, representing the point where the rate of decrease in the within-cluster sum of squares (WCSS) significantly slowed across network pairs, forming an “elbow” (cutoff = 0.95) (elbow for k varied between 28 and 34 across network pairs). This methodology ensures a balance between maximizing the cohesion within clusters and minimizing the total number of clusters.
  d. **K-means Cluster Affinity Metrics:** For each k-means cluster, we calculated the following subject level affinity metrics.
    i. Occupancy (OC): Cluster occupancy for a specific subject defined as the proportion of that subject’s total windows contained within the cluster, quantified the average duration a subject spent in that specific state of interaction between the network pairs.
    ii. Dwell time (DT): Dwell time for a subject and a cluster defined as the number of consecutive windows on average that fell within the cluster before moving out of the cluster, measured the average duration of a single, continuous instance of a network interaction state.
    iii. Distance to centroid - mean (DM) and std. (DSD): For each subject window, the Euclidean distance (d) to the cluster centroid is calculated. The mean and std. of this distance are then computed across all windows for a given subject. A smaller mean distance indicated a stronger subject affinity for the cluster interaction state across their entire trajectory, while a lower standard deviation suggested a more stable relationship.
    iv. Cluster similarity score (CSS) - mean (CSSM) and std. (CSSSD): CSS presents a complementary view on the k-means distances, where higher value means higher affinity. It employs a Gaussian kernel or radial basis function (RBF) kernel to measure the similarity between data points based on Euclidean distance (d) from a subject window to cluster centroid as given by:

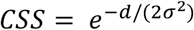

where σ is a smoothing parameter (we used median distance between all windows and their nearest cluster centroids). The subject level CSS mean across windows (CSSM) gives the subject’s average affinity to the cluster’s network interaction pattern across their entire trajectory, while std. (CSSSD) depicts the consistency or stability of this relationship.
5. **Statistical Evaluation** CN vs. SZ group differences in subject level scalar and cluster affinity metrics were assessed using robust regression with age, sex, and site as covariates. Multiple comparisons were corrected using a 5% False Discovery Rate (FDR) [21]. Robust linear regression analysis was also conducted to examine the relationship between subject-level joint density distribution metrics and several clinical measures with age, sex, and site as covariates. These measures included cognitive scores (for visual learning, verbal learning, processing speed, and working memory), symptom scores (PANSS Total), and drug scores (Chlorpromazine - CPZ - equivalent). For a more detailed explanation of these scores and the observed associations, please refer to [8].

**Figure 1.**
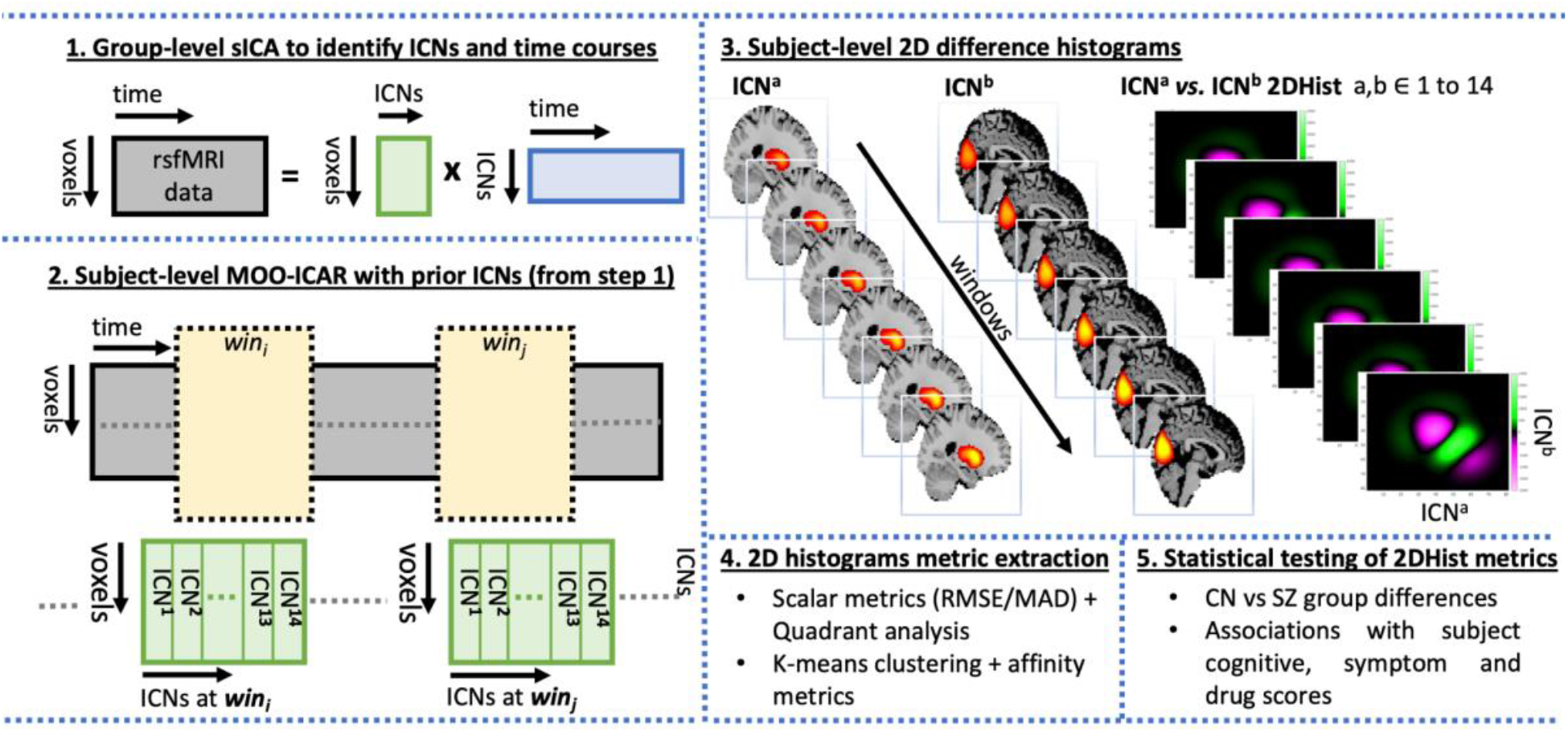
The DESINE Framework: The analysis pipeline proceeds through five main stages: (1) Group-level sICA to define the reference intrinsic connectivity networks (ICNs) and their associated time courses (2) Subject-level multi–objective optimization ICA with reference (MOO-ICAR), which uses prior ICNs to estimate spatially dynamic networks across sliding windows (3) Interaction Analysis where joint density distributions (2D Histograms) are generated for every network pair across windows, along with 2D difference histograms after subtracting the overall group mean joint histogram for a network pair from the window level data (4) Extraction of scalar metrics (RMSE/MAD), quadrant analysis, and their subject level means/standard deviations from the difference histograms, as well as k-means clustering and cluster affinity metrics (5) Statistical evaluation employing robust regression with age, sex and site as covariates to find significant group differences in various metrics between healthy controls (CN) and patients with schizophrenia (SZ), as well the associations of these metrics with subject cognitive, symptom and drug scores. See Section 2.2 for detailed description of the pipeline.

## 3. Results

### 3.1 Network Activity Maps and Interactions

The first two stages of the analysis pipeline from Section 2.2 and Figure 1 identified 14 spatially dynamic ICNs. The mean ICN maps for the CN group shown in Figure 2 exhibit a hierarchical organization of activity, where increasing the activity threshold progressively highlights focused, highly active core regions. The identified ICNs are: VIS-P (visual primary), SUB (subcortical), MTR-P (somatomotor primary), CER (cerebellar), ATN (attention-dorsal), FRNT (frontal), MTR-S (somatomotor secondary), FPN-R (frontoparietal right), VIS-S (visual secondary), pDMN (posterior default mode), FPN-L (frontoparietal left), SN (salience), TEMP (temporal), and aDMN (anterior default mode). Figure 3 illustrates the transient spatial interactions for the network pair VIS-P/FPN-R. 2D difference histograms and corresponding cluster mean maps reveal significant dynamic variability, showing how network regions overlap and activities fluctuate relative to the group mean state.

**Figure 2.**
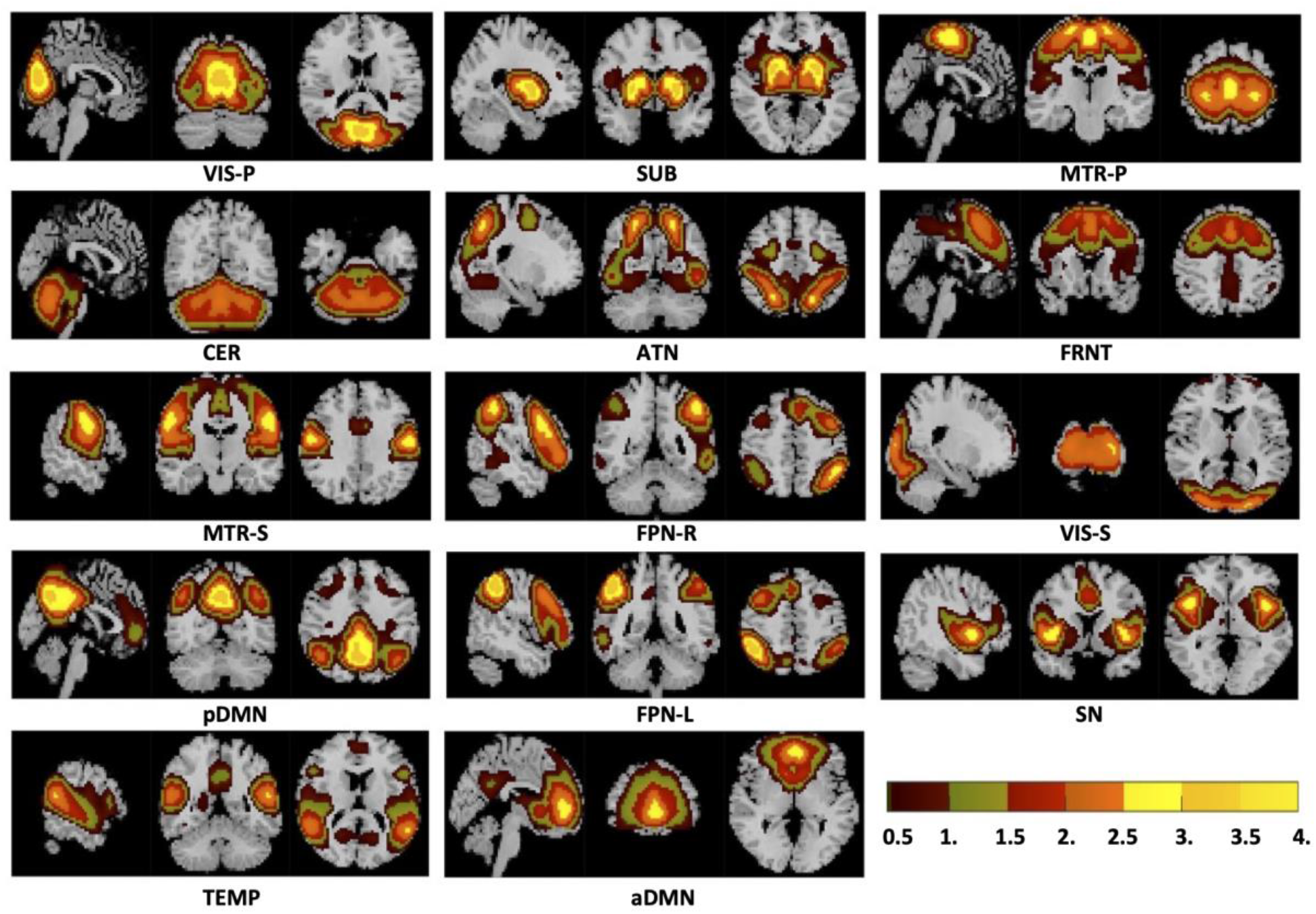
Network activity maps: Mean network activity calculated using z-scored data for the 14 spatially dynamic intrinsic connectivity networks (ICNs) in the control (CN) group shown. A threshold of *Z*_*th*_ = 0.5 employed alongside a discrete colormap progressively highlights the highly active core regions. The images are presented across three anatomical planes: sagittal, coronal, and transverse. The networks shown are: VIS-P (visual primary), SUB (subcortical), MTR-P (somatomotor primary), CER (cerebellar), ATN (attention-dorsal), FRNT (frontal), MTR-S (somatomotor secondary), FPN-R (frontoparietal right), VIS-S (visual secondary), pDMN (posterior default mode), FPN-L (frontoparietal left), SN (salience), TEMP (temporal), and aDMN (anterior default mode).

**Figure 3.**
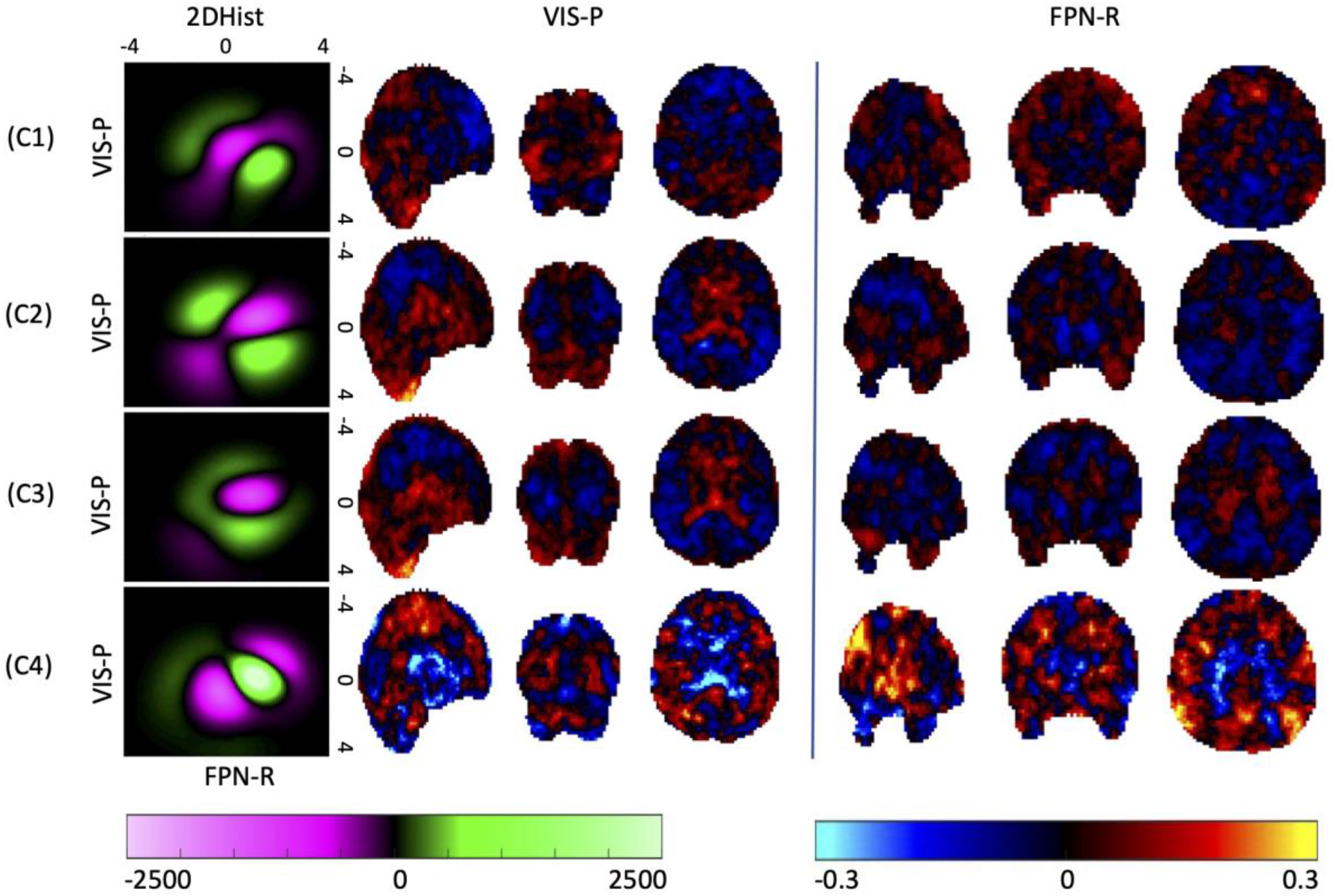
Cluster mean 2D histograms and corresponding network maps: Four different k-means clusters (C1-C4; k=35) and their mean 2D difference histograms are shown for the network pair VIS-P vs FPN-R. Normalized voxel-level activity for both the networks, ranging from -4 to +4 (mean at 0) at a resolution of size 0.1 is used to construct the 2D difference histograms of size 81×81 after subtracting the mean 2D histogram for the network pair. In addition, the corresponding mean activity maps across these clusters for both the networks are also shown across three anatomical planes (sagittal, coronal, and transverse) after subtracting the overall network mean map from the cluster mean map. Results highlight dynamical variability in network interactions across clusters.

### 3.2 Reduced Magnitude of Scalar Spatial Interaction Dynamics in Schizophrenia

CN vs. SZ group comparisons of scalar interaction metrics (per subject mean/std. of RMSE/MAD across windows) using robust regressions revealed a widespread and systematic reduction in the magnitude and variability of spatial interactions in SZ patients across several of the 91 possible network pairs, as summarized in Table 1 and the statistical heat maps in Figure 4,5, which illustrate the distribution of these differences across network pairs and quadrants.

**Table 1.** CN vs. SZ group differences in joint histogram metrics: Number of network pairs (out of total 91 combinations of 14 ICNs) showing significant CN vs. SZ group differences *(red, CN>SZ; blue, CN<SZ; black, total)* using robust regressions with age, sex and site as covariates after 5% FDR correction for multiple comparisons across network pairs. RMSE and MAD metrics for joint histograms of each network pair were computed at each subject-window and per subject mean/std. of these metrics across windows were used for analyzing group differences. Analysis was performed for either the whole joint histograms or separately for each of the four quadrants (Q1 - positive activity for both networks; Q2 - negative vs. positive; Q3 - negative vs. negative; Q4 - positive vs. negative) to separately analyze the interactions between the active and inactive regions of the network pairs. Results show that the metrics are predominantly higher in CN than SZ, indicating higher dynamic variability and interactions in the CN group. Only Q3 (with interactions between the inactive regions of both networks) shows one network pair with CN<SZ for std. metrics. Overall, Q1 with active regions of both networks shows the highest number of significant results. See Figure.4, 5 for heat maps revealing detailed network interactions.

| Region | Mean RMSE | Std RMSE | Mean MAD | Std MAD |
| --- | --- | --- | --- | --- |
| Whole | 47, 0, 47 | 0, 0, 0 | 17, 0, 17 | 0, 0, 0 |
| Q1 | 70, 0, 70 | 25, 0, 25 | 50, 0, 50 | 8, 0, 8 |
| Q2 | 19, 0, 19 | 10, 0, 10 | 8, 0, 8 | 2, 0, 2 |
| Q3 | 2, 0, 2 | 1, 1, 2 | 0, 0, 0 | 0, 1, 1 |
| Q4 | 19, 0, 19 | 5, 0, 5 | 13, 0, 13 | 0, 0, 0 |

**Figure 4.**
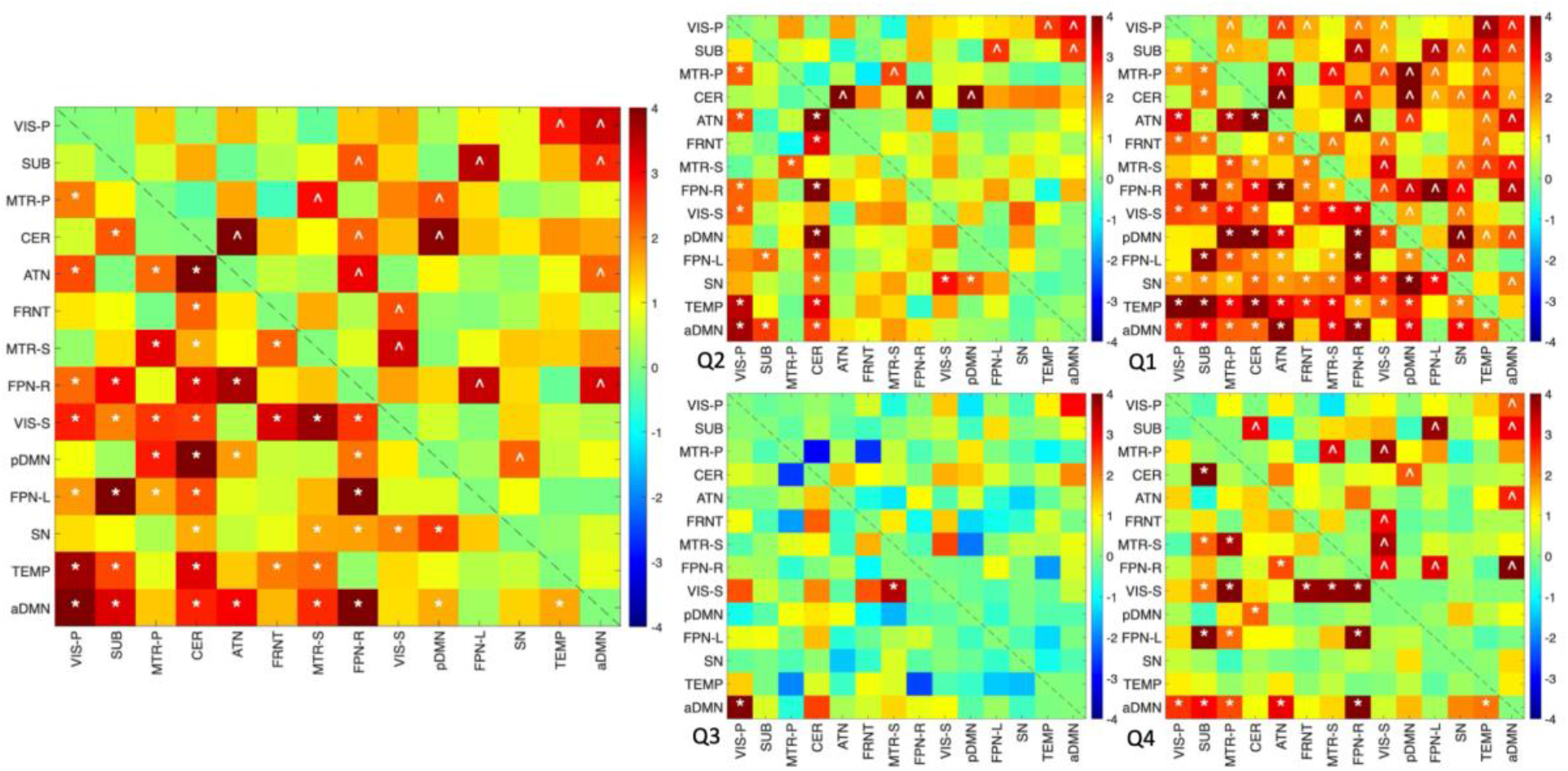
Heat maps of group differences in mean joint histogram metrics: CN vs. SZ group differences in subject wise mean RMSE (lower triangle matrix) and mean MAD (upper triangle matrix) computed using robust regressions with age, sex and site as covariates are shown. Colorbar represents the statistical values given by *k* = −*lo*g_10_(*p*) × *s*i*gn*(t). Results using entire joint histograms are shown on the left, while those computed separately for each quadrant (Q1-Q4) are shown on the right. Significant results after 5% FDR correction for multiple comparisons across network pairs are shown by the symbols * for mean RMSE and ^ for mean MAD. All significant results are positive (red) indicating higher values in CN than SZ, implying higher dynamic variability and interactions in the CN group. Q1 with active regions of both networks shows the highest number of significant results

**Figure 5.**
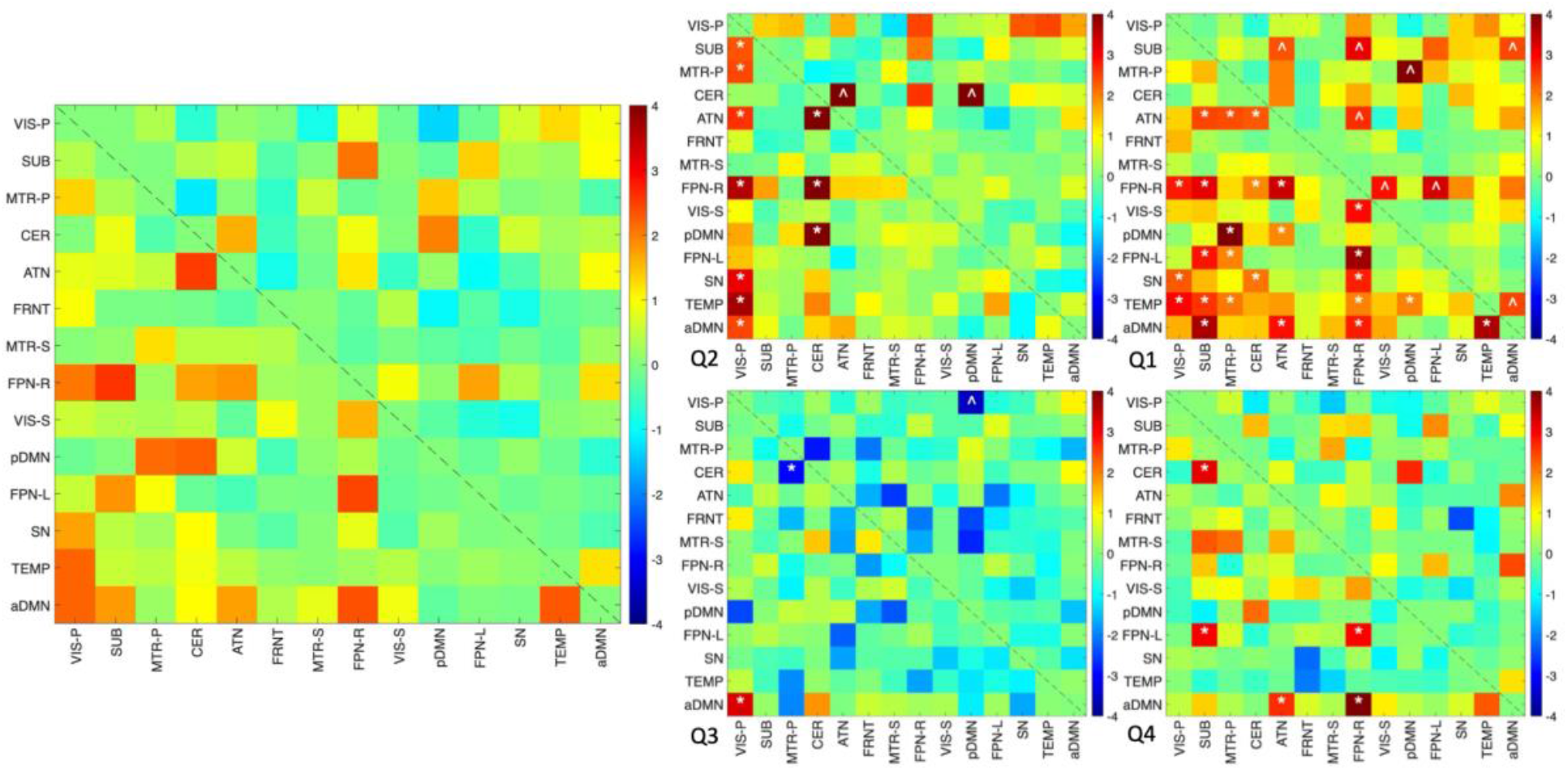
Heat maps of group differences in std. joint histogram metrics: CN vs. SZ group differences in subject wise std. RMSE (lower triangle matrix) and std. MAD (upper triangle matrix) computed using robust regressions with age, sex and site as covariates are shown. Colorbar represents the statistical values given by *k* = −*lo*g_10_(*p*) × *s*i*gn*(t). Results using entire joint histograms are shown on the left, while those computed separately for each quadrant (Q1-Q4) are shown on the right. Significant results after 5% FDR correction for multiple comparisons across network pairs are shown by the symbols * for std. RMSE and ^ for std. MAD. While the overall joint histograms analysis (left) doesn’t show any significant results, Q1 with active regions of both networks shows the highest number of significant results. Most of the significant results are positive (red) indicating higher values in CN than SZ, implying higher dynamic variability and interactions in the CN group. Only Q3 (with interactions between inactive regions of both networks) shows one network pair each with CN<SZ for std. RMSE and std. MAD.

For whole joint histograms, 47 network pairs showed significantly higher mean RMSE in CN group, and 17 pairs showed significantly higher mean MAD (CN > SZ) (see Table 1 and Figure 4). Crucially, no network pairs exhibited significantly higher interaction magnitude in the SZ group (CN < SZ), indicating a unidirectional deficiency in the patients’ interaction repertoire. To further localize these disruptions, we performed quadrant analysis to separate interactions between active regions (Q1: positive activity for both networks), inactive regions (Q3: negative activity for both), and mixed active-inactive regions (Q2/Q4). Figure 4 shows significant differences in subject-wise mean RMSE (lower triangle) and mean MAD (upper triangle) visualized using a mapping of the statistic *k* = − log_10_(*p*) ∗ *s*i*gn*(t), where t is the ratio of the regression coefficient to the standard error of its estimate. For mean metrics, significant results after 5% FDR correction were found exclusively in the positive range (red), confirming higher dynamic variability in CN than SZ. The highest density of significant differences occurred in Quadrant 1, where 70 network pairs (RMSE) and 50 pairs (MAD) showed CN > SZ effects, highlighting that interactions between the highly active core regions of networks are the most severely constrained in schizophrenia, while other regions also show dysfunction to a lesser extent.

Analysis of the standard deviation (std.) of these metrics across windows provided insights into the flexibility of spatial interactions (Table 1 and heat maps in Figure 5). While analysis of the entire joint histograms showed no significant group differences in variability, the quadrant-specific analysis revealed significant findings, predominantly in Q1. 25 pairs showed significantly higher std. RMSE and 8 pairs showed higher std. MAD in controls in Q1, suggesting that healthy individuals express a more fluid and flexible range of spatial interaction. Conversely, SZ patients exhibit a more “rigid” or constrained variability in these active-active interactions. Interestingly, Q3 (interactions between inactive regions) provided the only exception to the CN > SZ trend, showing one network pair each where SZ patients exhibited significantly higher variability than controls (CN < SZ) for std. RMSE (CER/MTR-P) and std. MAD (VIS-P/pDMN). Overall, the results suggest “spatial rigidity” observed in SZ is primarily localized to the interactions of active network regions.

### 3.3 Cognitive and Clinical Associations of Scalar Spatial Interaction Metrics

To investigate the functional relevance of the observed spatial dynamics, we evaluated the associations between the scalar metrics of whole joint histograms as well as separate quadrants and several clinical and cognitive subject scores. As detailed in Table 2, statistical significance was determined using robust regressions with age, sex, and site as covariates, followed by 5% FDR correction for multiple comparisons.

**Table 2.**
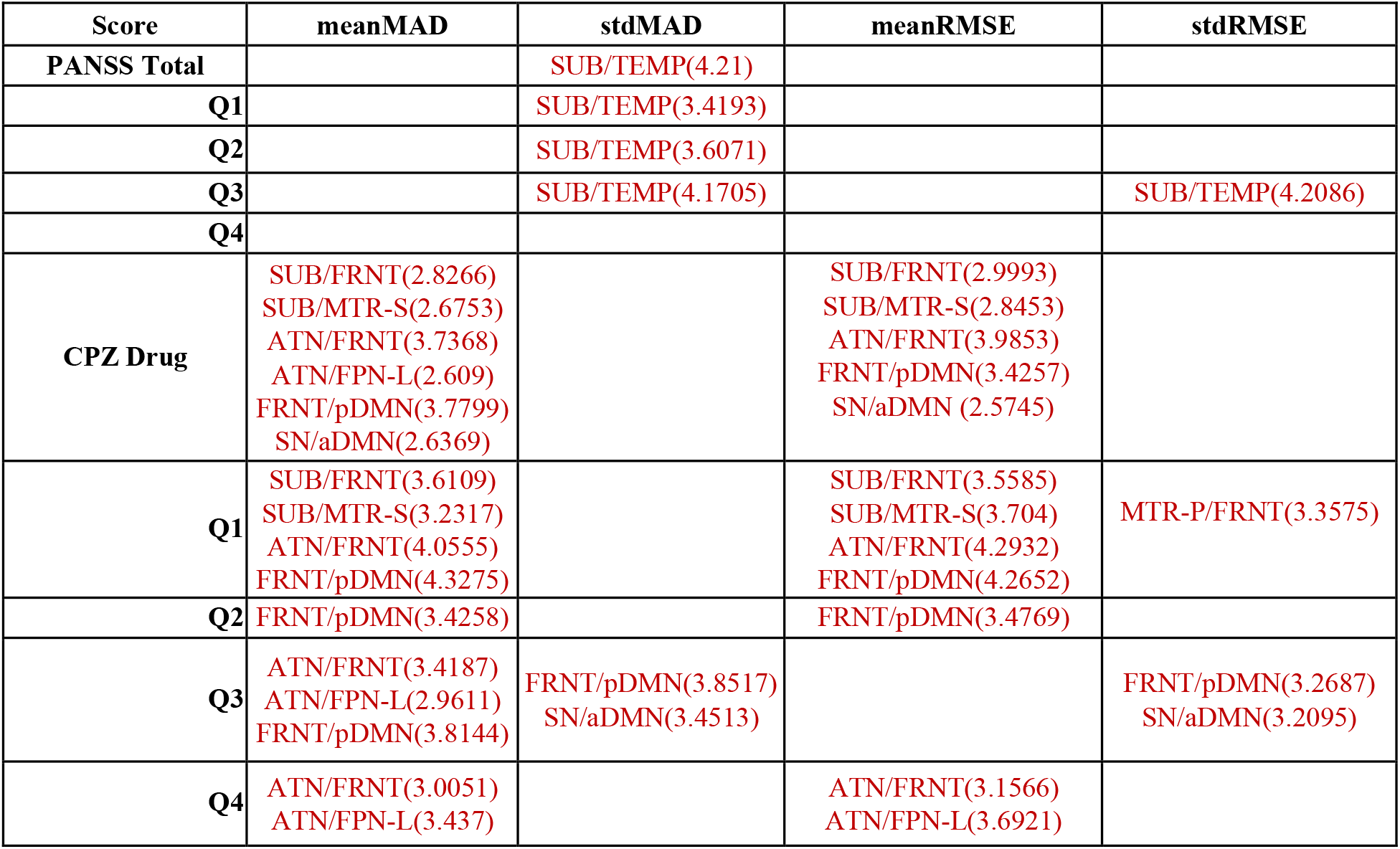
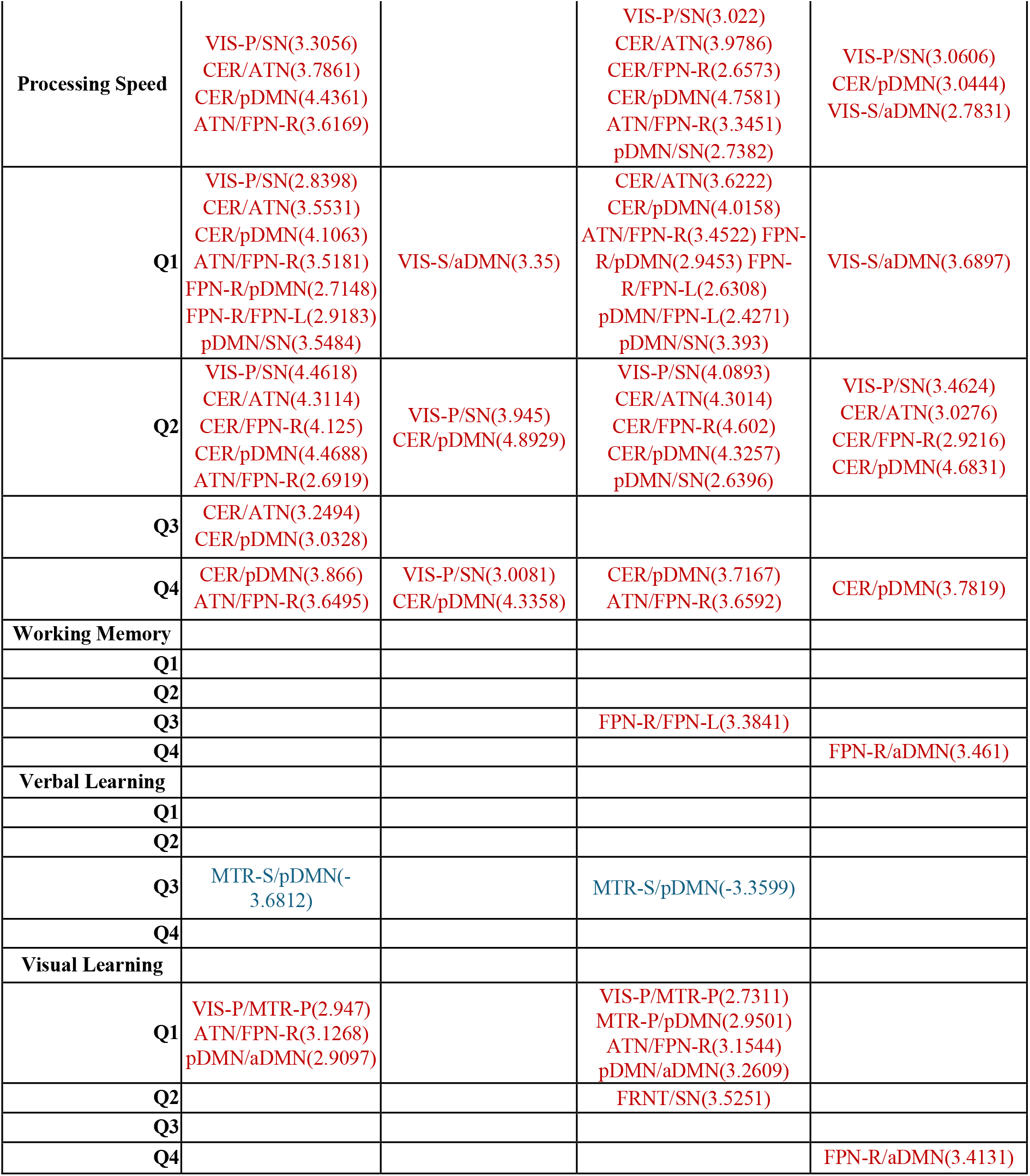
Cognitive, Clinical and Drug Score Associations of Scalar Spatial Interaction Metrics: For each score, the table lists the network pairs with significant association with subject level mean/std. of RMSE/MAD metrics, after 5% FDR correction for multiple comparisons across network pairs. Results using full joint histograms for a score are shown in the first row, followed by results for each quadrant of the joint histograms in a separate row below. All values shown are *k* = −*lo*g_10_(*p*) × *s*i*gn*(t), where t is the ratio of the regression coefficient to the standard error of its estimate. Positive associations are shown in red, while negative ones are shown in blue. The associations are predominantly positive (red), indicating higher dynamic variability and interactions result in higher scores. Only verbal learning shows negative association (blue) with mean RMSE/MAD for the network pair MTR-S/pDMN in Q3 (interaction between non-active regions of both networks).

| Score | meanMAD | stdMAD | meanRMSE | stdRMSE |
| --- | --- | --- | --- | --- |
| PANSS Total |  | SUB/TEMP(4.21) |  |  |
| Q1 |  | SUB/TEMP(3.4193) |  |  |
| Q2 |  | SUB/TEMP(3.6071) |  |  |
| Q3 |  | SUB/TEMP(4.1705) |  | SUB/TEMP(4.2086) |
| Q4 |  |  |  |  |
| CPZ Drug | SUB/FRNT(2.8266)<br>SUB/MTR-S(2.6753)<br>ATN/FRNT(3.7368)<br>ATN/FPN-L(2.609)<br>FRNT/pDMN(3.7799)<br>SN/aDMN(2.6369) |  | SUB/FRNT(2.9993)<br>SUB/MTR-S(2.8453)<br>ATN/FRNT(3.9853)<br>FRNT/pDMN(3.4257)<br>SN/aDMN (2.5745) |  |
| Q1 | SUB/FRNT(3.6109)<br>SUB/MTR-S(3.2317)<br>ATN/FRNT(4.0555)<br>FRNT/pDMN(4.3275) |  | SUB/FRNT(3.5585)<br>SUB/MTR-S(3.704)<br>ATN/FRNT(4.2932)<br>FRNT/pDMN(4.2652) | MTR-P/FRNT(3.3575) |
| Q2 | FRNT/pDMN(3.4258) |  | FRNT/pDMN(3.4769) |  |
| Q3 | ATN/FRNT(3.4187)<br>ATN/FPN-L(2.9611)<br>FRNT/pDMN(3.8144) | FRNT/pDMN(3.8517)<br>SN/aDMN(3.4513) |  | FRNT/pDMN(3.2687)<br>SN/aDMN(3.2095) |
| Q4 | ATN/FRNT(3.0051)<br>ATN/FPN-L(3.437) |  | ATN/FRNT(3.1566)<br>ATN/FPN-L(3.6921) |  |
| Processing Speed | VIS-P/SN(3.3056)<br>CER/ATN(3.7861)<br>CER/pDMN(4.4361)<br>ATN/FPN-R(3.6169) |  | VIS-P/SN(3.022)<br>CER/ATN(3.9786)<br>CER/FPN-R(2.6573)<br>CER/pDMN(4.7581)<br>ATN/FPN-R(3.3451)<br>pDMN/SN(2.7382) | VIS-P/SN(3.0606)<br>CER/pDMN(3.0444)<br>VIS-S/aDMN(2.7831) |
| Q1 | VIS-P/SN(2.8398)<br>CER/ATN(3.5531)<br>CER/pDMN(4.1063)<br>ATN/FPN-R(3.5181)<br>FPN-R/pDMN(2.7148)<br>FPN-R/FPN-L(2.9183)<br>pDMN/SN(3.5484) | VIS-S/aDMN(3.35) | CER/ATN(3.6222)<br>CER/pDMN(4.0158)<br>ATN/FPN-R(3.4522) FPN-R/pDMN(2.9453) FPN-R/FPN-L(2.6308)<br>pDMN/FPN-L(2.4271)<br>pDMN/SN(3.393) | VIS-S/aDMN(3.6897) |
| Q2 | VIS-P/SN(4.4618)<br>CER/ATN(4.3114)<br>CER/FPN-R(4.125)<br>CER/pDMN(4.4688)<br>ATN/FPN-R(2.6919) | VIS-P/SN(3.945)<br>CER/pDMN(4.8929) | VIS-P/SN(4.0893)<br>CER/ATN(4.3014)<br>CER/FPN-R(4.602)<br>CER/pDMN(4.3257)<br>pDMN/SN(2.6396) | VIS-P/SN(3.4624)<br>CER/ATN(3.0276)<br>CER/FPN-R(2.9216)<br>CER/pDMN(4.6831) |
| Q3 | CER/ATN(3.2494)<br>CER/pDMN(3.0328) |  |  |  |
| Q4 | CER/pDMN(3.866)<br>ATN/FPN-R(3.6495) | VIS-P/SN(3.0081)<br>CER/pDMN(4.3358) | CER/pDMN(3.7167)<br>ATN/FPN-R(3.6592) | CER/pDMN(3.7819) |
| Working Memory |  |  |  |  |
| Q1 |  |  |  |  |
| Q2 |  |  |  |  |
| Q3 |  |  | FPN-R/FPN-L(3.3841) |  |
| Q4 |  |  |  | FPN-R/aDMN(3.461) |
| Verbal Learning |  |  |  |  |
| Q1 |  |  |  |  |
| Q2 |  |  |  |  |
| Q3 | MTR-S/pDMN(-3.6812) |  | MTR-S/pDMN(-3.3599) |  |
| Q4 |  |  |  |  |
| Visual Learning |  |  |  |  |
| Q1 | VIS-P/MTR-P(2.947)<br>ATN/FPN-R(3.1268)<br>pDMN/aDMN(2.9097) |  | VIS-P/MTR-P(2.7311)<br>MTR-P/pDMN(2.9501)<br>ATN/FPN-R(3.1544)<br>pDMN/aDMN(3.2609) |  |
| Q2 |  |  | FRNT/SN(3.5251) |  |
| Q3 |  |  |  |  |
| Q4 |  |  |  | FPN-R/aDMN(3.4131) |

#### Cognitive associations

The associations were found to be predominantly positive (red in Table 2), suggesting that individuals who exhibit higher dynamic variability and magnitude of spatial interactions also tend to have higher cognitive and clinical scores. Specifically, processing speed and visual learning showed robust positive associations with scalar spatial interaction metrics across multiple network pairs and regions. For instance, higher mean MAD/RMSE for CER/pDMN across regions was positively linked to superior processing speed. Interactions between visual and motor systems (VIS-P/MTR-P) and attentional networks (ATN/FPN-R) were associated with better visual learning performance. Working memory was positively associated with RMSE metrics for FPN-R/FPN-L and FPN-R/aDMN networks.

#### Clinical and drug associations

Interaction metrics were positively associated with CPZ equivalent drug scores and PANSS symptom scores. Significant associations with CPZ scores were observed in network pairs such as ATN/FRNT and FRNT/pDMN, while PANSS total scores were linked to interaction flexibility (std. metrics) in SUB/TEMP. These findings suggest that both medication levels and symptom severity are associated with the expressed magnitude and variability of the “spatial dynamic repertoire.”

An exception to this positive trend was observed for verbal learning, which showed a significant negative association (blue in Table 2) with mean RMSE and MAD for the MTR-S/pDMN pair, specifically in quadrant 3. This indicates that in the inactive or non-core regions of these networks, higher interaction magnitude is associated with lower verbal learning scores. Aside from this localized finding in inactive regions, the global pattern suggests that the inability to express high-magnitude and diverse spatial interactions as seen in Section 3.2 (Table 1, Figure 4,5) contributes directly to the cognitive deficits characteristic of Schizophrenia.

### 3.4 Altered Dynamic Interaction State Affinities in Schizophrenia

To characterize the recurring patterns of spatial interactions, we applied k-means clustering (with k=35) to 2D difference histograms across 91 network pairs. As summarized in Table 3, robust regressions revealed several significant CN vs. SZ group differences across six cluster affinity metrics: occupancy (OC), dwell time (DT), distance means (DM), distance standard deviation (DSD), cluster similarity score mean (CSSM), and cluster similarity score standard deviation (CSSSD) with 5% FDR correction for multiple comparisons (3185 total comparisons for each metric).

**Table 3.** Group differences in k-means cluster metrics of joint histograms: Number of clusters with significant CN vs. SZ group differences in 6 different metrics of k-means clusters (k=35) using robust regressions with age, sex and site as covariates, after 5% FDR correction for multiple comparisons across network pairs and clusters (35 clusters * 91 network pairs = 3185 comparisons). The number of significant results with CN>SZ are shown in red, CN<SZ are shown in blue, and the total in black. See Figure.6 for results highlighted for a few clusters for the network pair VIS-P vs FPN-R as depicted in Figure.3, and see Figure.7 for connectograms highlighting the most significant network pair results.

|  | Occupancy | Dwell time | Distance means | Distance std. | CSS Means | CSS std |
| --- | --- | --- | --- | --- | --- | --- |
| **CN > SZ** | 291 | 113 | 998 | 264 | 366 | 384 |
| **CN < SZ** | 292 | 99 | 344 | 20 | 942 | 700 |
| **Total** | 583 | 212 | 1342 | 284 | 1308 | 1084 |

The analysis revealed significant group differences, with metrics primarily highlighting altered affinity in patients for standard interaction states. Specifically, we identified 583 significant clusters for occupancy and 212 for dwell time, revealing how patients and controls spend different amounts of time in various interaction states/clusters. Distance-based metrics showed widespread disruptions, with over 1,300 clusters exhibiting significant differences in DM and CSSM, 284 clusters in DSD and 1084 clusters in CSSSD. As shown in Table 3, most of these results (CN > SZ for DM/DSD and CN < SZ for CSSM/CSSSD) indicate that SZ patients show lower variability, lower flexibility or higher rigidity in their dynamic repertoire and their affinities towards various interaction states.

Group differences for four example clusters for the VIS-P/FPN-R network pair are highlighted in Figure 6. SZ patients exhibited significantly higher occupancy and dwell time in certain interaction states (clusters 2/3), suggesting an atypical persistence in specific spatial configurations. Conversely, they showed significantly lower DM and higher CSSM in clusters 1/2/3, indicating lower flexibility and increased rigidity in their spatial structures and interactions when they enter a particular functional state.

**Figure 6.**
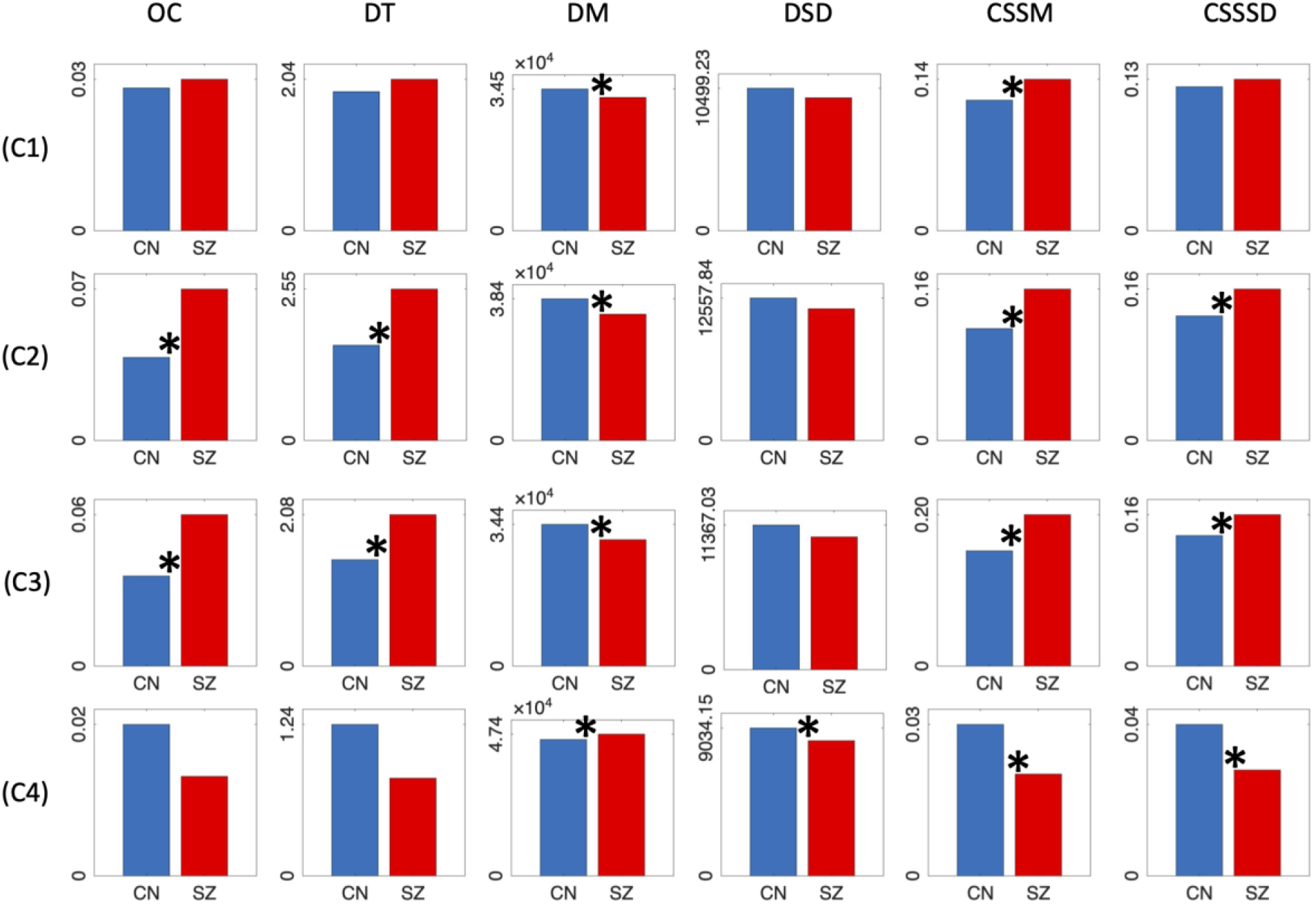
Group differences in k-means cluster metrics of joint histograms highlighted for VIS-P/FPN-R: CN vs. SZ group differences in cluster metrics - occupancy (OC), dwell time (DT), distance means (DM), distance std. (DSD), CSS means (CSSM), and CSS std. (CSSSD) - are shown for four clusters for the network pair VIS-P/FPN-R, whose joint histograms and mean network maps are depicted in Figure.3. The symbol * highlights significant group differences after 5% FDR correction for multiple comparisons across network pairs and clusters (35 clusters * 91 network pairs = 3185 comparisons) for each metric.

The sheer scale of these disruptions across six metrics is visualized in the multicluster connectograms of Figure 7. Given that a 5% FDR threshold for significance would result in nearly fully connected graphs (Table 3), we employed modified higher thresholds to isolate the top 10% of significant disruptions (e.g., the top 58 results for occupancy out of 583 total results). These connectograms illustrate that spatial interaction deficits in schizophrenia are not localized to a single circuit but represent a global breakdown in the brain’s ability to maintain high-fidelity, diverse spatial configurations.

**Figure 7.**
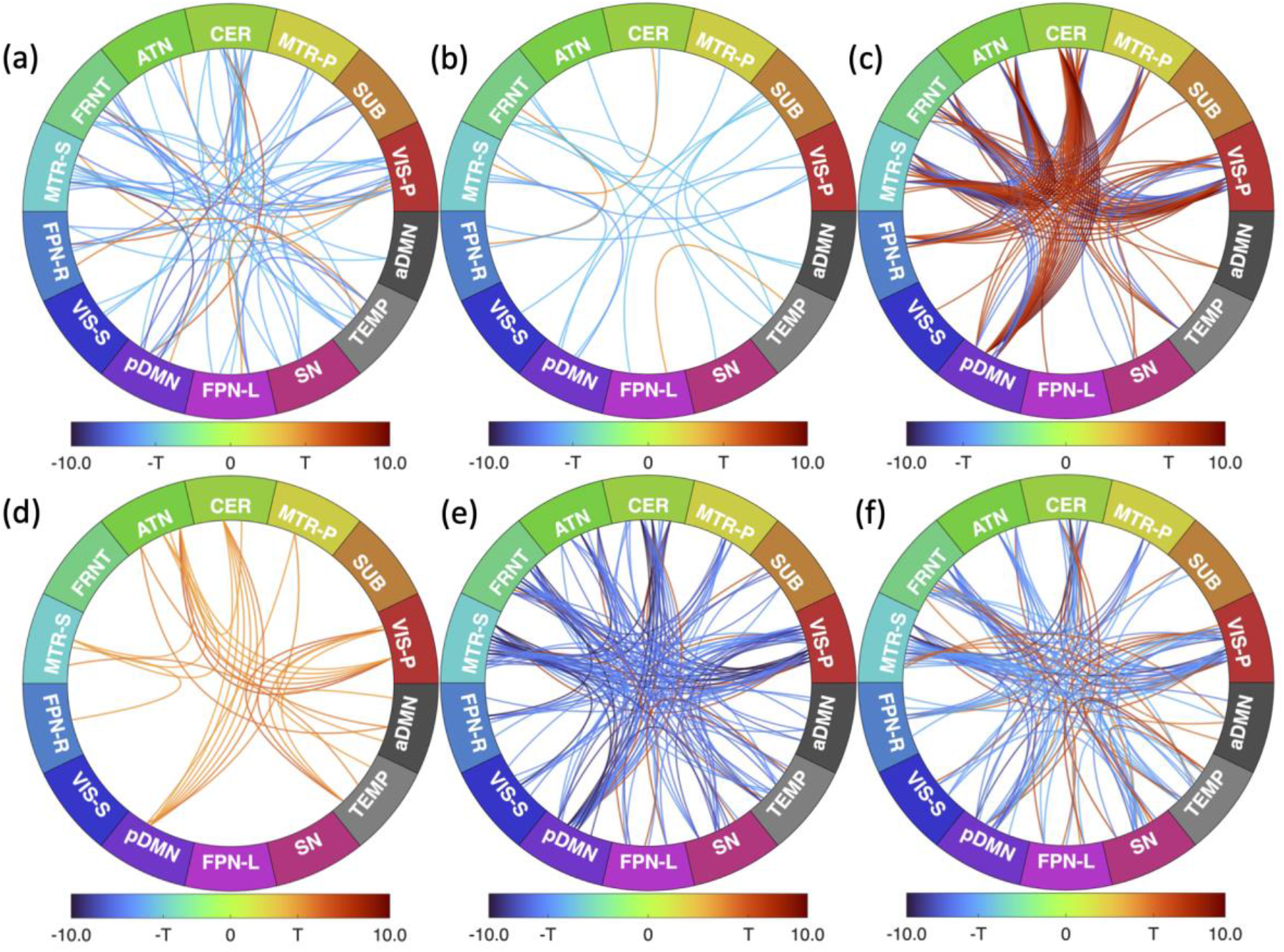
Multicluster-connectograms highlighting the most significant CN vs. SZ group differences in k-means cluster metrics of joint histograms: The most significant results for the network pairs from Table.3 are highlighted here for the six metrics of k-means clusters of 2D difference histograms: (a) occupancy, (b) dwell time, (c) distance means, (d) distance std., (e) CSS means, and (f) CSS std. Multiple connections between a network pair highlight multiple significant clusters, with red connections showing CN>SZ and blue CN<SZ. Colorbars and connections represent the statistic value *k* = − log_10_(*p*) ∗ *s*i*gn*(t). A threshold of *k*_*cr*_ = − log_10_(*p*_*cr*_), where *p*_*cr*_ represents the critical p-value for significance after 5% FDR correction, would highlight all the significant connections with *abs*(*k*) ≥ *k*_*cr*_ . This value for each of the metric is: (a) 2.0389 (b) 2.4906 (c) 1.6835 (d) 2.3525 (e) 1.6893 (f) 1.7696. Since this would result in a fully connected graph for each metric due to the plethora of significant results, we only highlighted the top 10% of the significant results with *abs*(*k*) ≥ *T*, where T represents a modified threshold for each metric chosen to depict only the top 10% most prominent results, after 5% FDR correction. For example (a) highlights 58 prominent results (top 10%) out of the 583 total significant results for occupancy (after FDR) from Table 3.

### 3.5 Cognitive and Clinical Associations of Dynamic Interaction State Affinities

We found that the affinities for specific dynamic spatial interaction states are strongly linked to clinical, drug and cognitive outcomes, as summarized in Table 4. We evaluated associations across 3185 comparisons (35 clusters × 91 network pairs) for each score-metric combination using robust regression with 5% FDR correction. Detailed results for score-metric combinations (Figure 8,9) highlight a wide range of associations varying from sparse to dense and from highly negative to highly positive.

**Table 4.** Cognitive, Clinical and Drug Score Associations of k-means cluster metrics of joint histograms: No of clusters showing significant associations under robust regression with age, sex, site as covariates, after 5% FDR correction for multiple comparisons (35 clusters * 91 network pairs = 3185 comparisons) for each score vs. metric combination are shown. The number of *positive, negative* and *total* significant associations are shown in red, blue and black respectively. See Figure.8,9 for detailed results of significant associations between subject scores and k-means metrics.

|  | Occupancy | Dwell time | Distance means | Distance std. | CSS Means | CSS std. |
| --- | --- | --- | --- | --- | --- | --- |
| **PANSS total** | 0, 0, 0 | 0, 0, 0 | 0, 0, 0 | 4, 0, 4 | 0, 0, 0 | 0, 0, 0 |
| **CPZ drug** | 39, 0, 39 | 23, 0, 23 | 3, 0, 3 | 0, 0, 0 | 6, 0, 6 | 2, 0, 2 |
| **Processing Speed** | 4, 20, 24 | 2, 2, 4 | 344, 108, 452 | 86, 0, 86 | 44, 241, 285 | 43, 72, 115 |
| **Working Memory** | 1, 0, 1 | 0, 0, 0 | 0, 0, 0 | 0, 0, 0 | 0, 0, 0 | 2, 1, 3 |
| **Verbal Learning** | 6, 0, 6 | 0, 0, 0 | 55, 59, 114 | 0, 0, 0 | 22, 9, 31 | 20, 10, 30 |
| **Visual Learning** | 10, 23, 33 | 1, 7, 8 | 383, 156, 539 | 5, 0, 5 | 53, 307, 360 | 50, 169, 219 |

**Figure 8.**
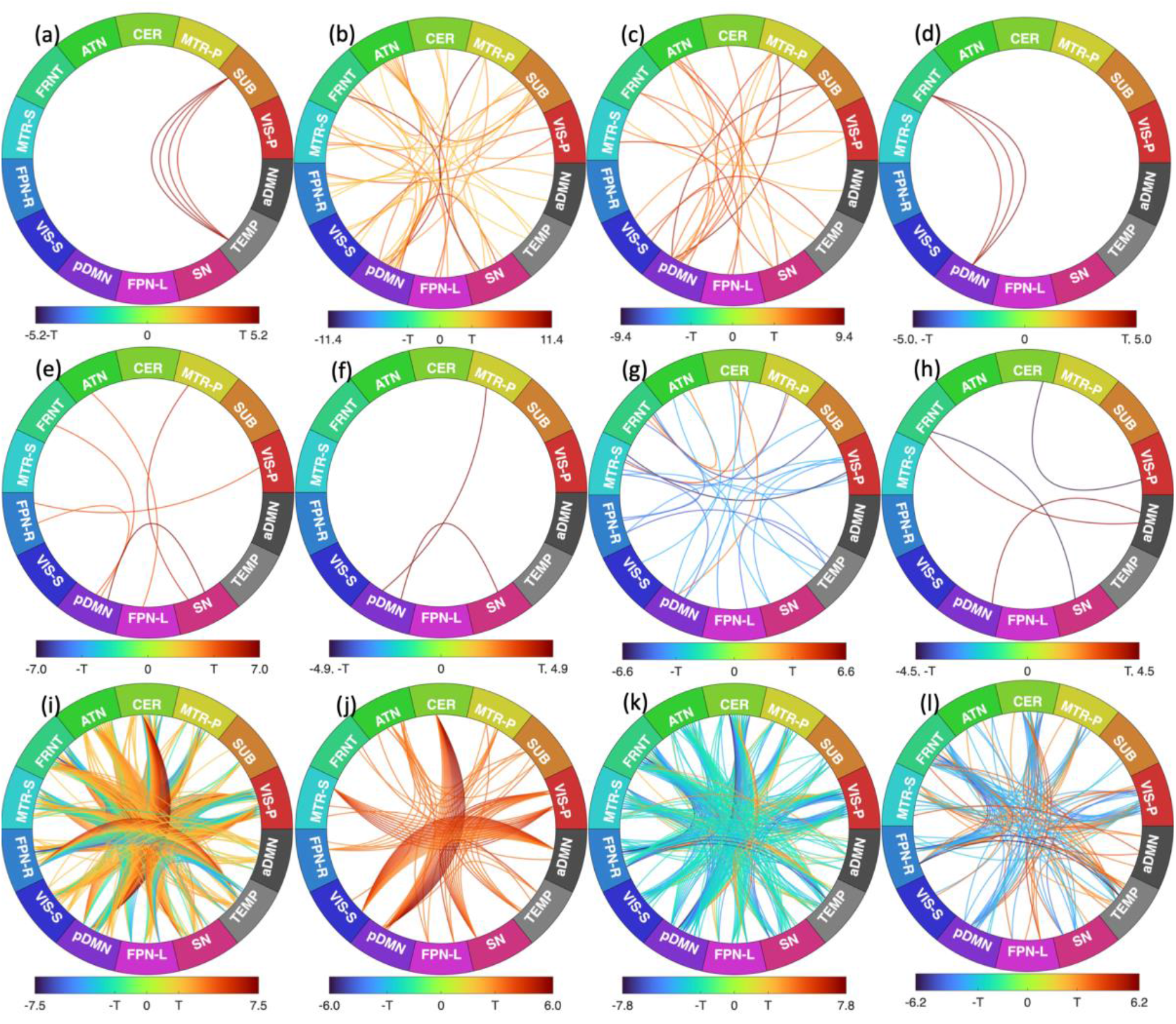
Multicluster-connectograms showing the association of subject symptom, drug, cognitive scores with k-means cluster metrics of joint histograms: Significant network-pair-clusters vs subject score associations are shown here for (a) PANSS Total vs. Distance std. (b) CPZ drug vs. occupancy (c) CPZ drug vs. dwell time (d) CPZ drug vs. distance means (e) CPZ drug vs. CSS means (f) CPZ drug vs. CSS std. (g) Processing Speed vs. occupancy (h) Processing Speed vs. dwell time (i) Processing Speed vs. distance means (j) Processing Speed vs. distance std. (k) Processing Speed vs. CSS means (l) Processing speed vs. CSS std. Multiple connections between a network pair highlight multiple significant clusters, with red connections showing positive associations and blue negative. Colorbars and connections represent the statistic value *k* = −*lo*g_10_(*p*) × *s*i*gn*(t). A threshold of *T* = *k*_*cr*_ = −*lo*g_10_(*p*_*cr*_), where p_cr_ represents the critical p-value for significance after 5% FDR correction is used to show all the significant connections with *abs*(*k*) ≥ *T* in each connectogram. The results indicate a broad spectrum of associations, ranging from sparse to dense and from strongly negative to highly positive.

**Figure 9.**
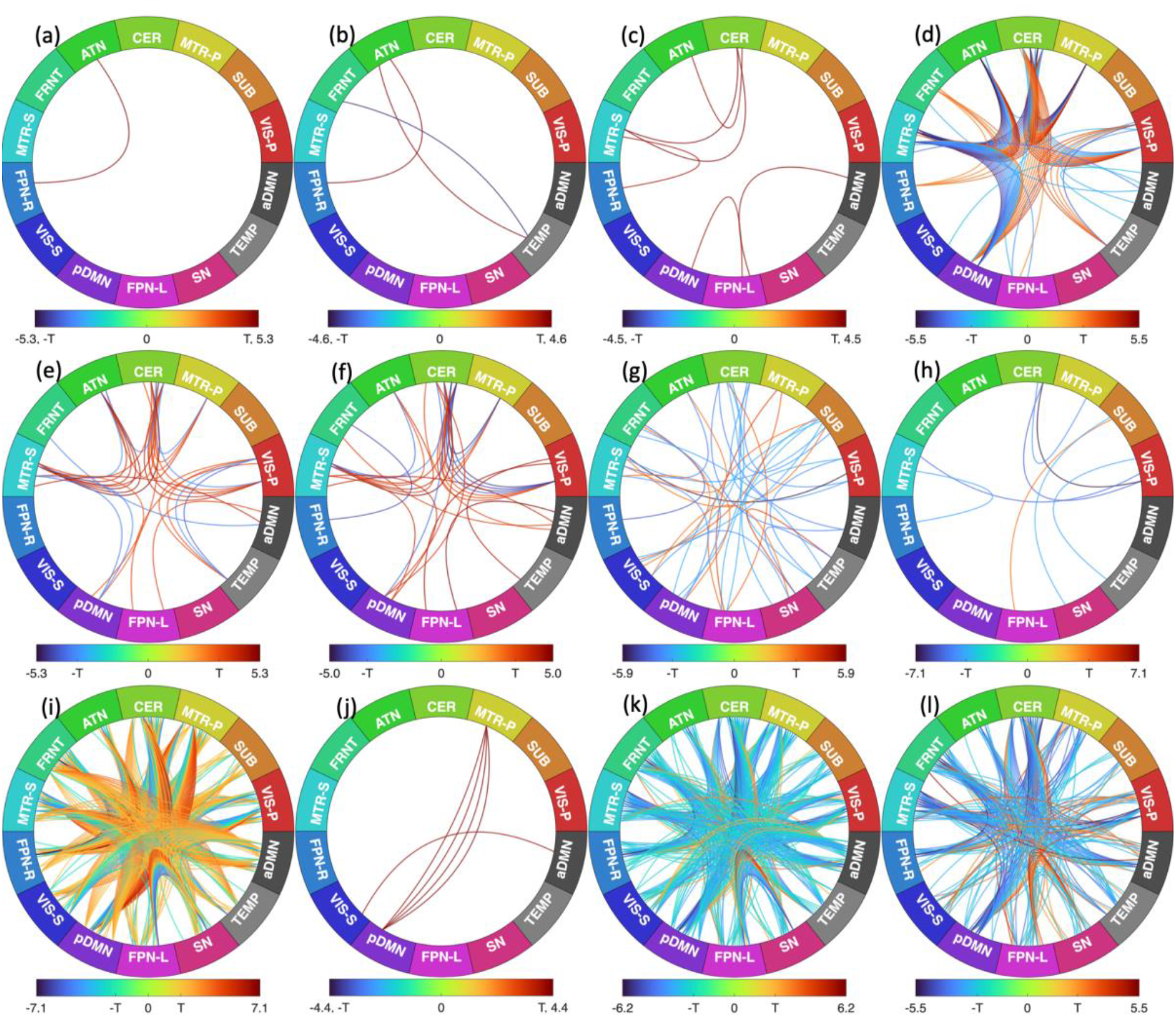
Multicluster-connectograms showing the association of subject symptom, drug, cognitive scores with k-means cluster metrics of joint histograms: Significant network-pair-clusters vs subject score associations are shown here for (a) Working memory vs. occupancy (b) Working memory vs. CSS std. (c) Verbal learning vs. occupancy (d) Verbal learning vs. distance means (e) Verbal learning vs. CSS means (f) Verbal learning vs. CSS std. (g) Visual learning vs. occupancy (h) Visual learning vs. dwell time (i) Visual learning vs. distance means (j) Visual learning vs. distance std. (k) Visual learning vs. CSS means (l) Visual learning vs. CSS std. All other details similar to Figure 8. The associations observed demonstrate a broad spectrum, ranging from low to high density and covering a wide range from strongly negative to strongly positive.

#### Cognitive Associations

Similar to the associations with scalar metrics (Table 2, Section 3.2), processing speed and visual learning showed the most robust associations with affinity metrics, followed by verbal learning and working memory. Higher (red) DM/DSD and lower (blue) CSSM/CSSD (as also predominantly seen in the control group in Table 3, Section 3.4) were largely associated with improved processing speeds and visual learning (Table 4, Figure. 8i-l, 9i-l), while a few clusters also showed opposite behaviors indicating that the brain’s ability to precisely express and maintain specific spatial interaction patterns is a critical substrate for diverse cognitive functions. For example, Figure 9d shows that the distance means for several pDMN/CER clusters were positively associated (red) with verbal learning while those of pDMN/MTR-S were negatively associated (red).

#### Clinical and Drug Associations

Dynamic interaction state affinity metrics were significantly associated with clinical status. CPZ equivalent drug scores showed consistently positive associations with state occupancy and dwell time, as well as distance-based metrics (eg. pDMN/FRNT, pDMN/SN). This indicates that antipsychotic medication doses may influence the spatiotemporal characteristics of network interactions. Furthermore, PANSS total scores were specifically linked to the stability of SUB/TEMP interaction states, showing significant associations with DSD. This suggests that more inconsistent adherence to spatial interaction states (higher DSD) is associated with increased symptom severity.

Overall, the widespread nature of these associations, particularly with cognitive domains like processing speed and visual learning, reinforces the hypothesis that spatial interaction rigidity and altered state affinity observed in schizophrenia are central to the cognitive and clinical dysfunction associated with the disorder.

## 4. Discussion and Conclusions

In this study, we introduced a joint density distribution framework to characterize time-varying spatial interactions of brain networks. Our results provide compelling evidence that schizophrenia is characterized by a “constrained dynamic repertoire” [8,9], where network interactions become rigid and systematically dampened.

### 4.1 Constrained Repertoire and Spatial Rigidity

The scalar analysis reveals a systematic reduction in mean RMSE and MAD metrics in SZ patients, quantifying a significant loss in the magnitude of transient deviation from the average functional interaction state. While healthy brains (CN) exhibit fluid, high-magnitude spatial reorganizations expanding into each other’s territory or creating complex configurations of spatial overlap, predominantly within active-active network regions (Quadrant 1), SZ network interactions appear “stuck” closer to the group mean. This “spatial rigidity” in schizophrenia aligns with previous findings of reduced volumetric variability [8] and a synergistic loss of dynamic flexibility in spatial complexity [9], suggesting that the inability to express diverse, high-magnitude spatial configurations is a fundamental feature of the disorder’s psychopathology.

### 4.2 Altered State Affinity and Loss of Fidelity

The k-means cluster analysis revealed a global breakdown in the brain’s ability to maintain high-fidelity spatial configurations, with SZ patients exhibiting lower distance means (DM) and higher cluster similarity score means (CSSM) consistent with increased rigidity or loss of flexibility in interaction patterns. Additionally, patients spent atypical amounts of time (occupancy/dwell time) in certain interaction states, suggesting the SZ brain is prone to becoming “trapped” in altered configurations, which may provide a potential mechanism for the observed cognitive impairment [22].

### 4.3 Functional Relevance and Clinical Implications

Results link spatial interaction dynamics to clinical, drug, and cognitive indicators. Positive association between processing speed/learning and interaction metrics (RMSE/MAD/affinity) suggest that diverse spatial configurations are essential for efficient information processing. Interaction rigidity likely contributes to cognitive impairment. Symptom severity (PANSS Total) was tied to interaction stability in SUB/TEMP (std. RMSE, std. MAD, distance standard deviation), suggesting inconsistency in their interaction patterns linked to increased symptoms. CPZ equivalent drug scores were positively associated with scalar and affinity metrics, implying medication might influence spatio-temporal characteristics to compensate for rigidity.

### 4.4 Conclusion

This study demonstrates that the spatial interactions of brain networks at rest are significantly altered in schizophrenia. By utilizing DESINE, a global, voxel-agnostic 2D histogram framework, we identified a pervasive reduction in the magnitude, flexibility, and fidelity of these interactions. Our results link this “constrained dynamic repertoire” to cognitive deficits and clinical symptoms, positioning spatial interaction rigidity as a fundamental feature of the schizophrenia brain. This framework provides a novel, quantifiable biomarker for understanding network-level dysfunction in complex psychiatric and neurological disorders.

## Acknowledgments

This work was supported in part by the NSF grant 2112455 and the NIH grant R01MH123610, both awarded to Dr. Vince D. Calhoun, as well as the NIH grant 5R01MH119251, awarded to Dr. Armin Iraji. We gratefully acknowledge the members of the TReNDS Center for their valuable discussions.

## Disclosure

The authors declare no conflicts of interest.

## Notes

### Competing Interest Statement

The authors have declared no competing interest.

